# Dynamic coupling of cell fate specification and cell sorting during mouse preimplantation development

**DOI:** 10.64898/2026.09.01.748504

**Authors:** Sascha Ollertz, Silvia Muñoz-Descalzo, Sabine C. Fischer

**Affiliations:** Julius-Maximilians-Universität Würzburg,Faculty of Biology, CAIDAS, Biocenter, WueBiT, Chair for Computational and Theoretical Biology (CCTB), Klara-Oppenheimer-Weg 32, 97074 Würzburg, Germany; Instituto Universitario de Investigaciones Biomédicas y Sanitarias (IUIBS), Universidad Las Palmas de Gran Canaria (ULPGC), Paseo Blas Cabrera Felipe “Físico” 17, Las Palmas de Gran Canaria 35016, Spain

**Keywords:** Pattern formation, Cell differentiation, Mathematical Modelling, Differential Adhesion

## Abstract

During preimplantation development in mice, cells of the inner cell mass undergo a cell fate decision to become either Epiblast (Epi) or Primitive Endoderm (PrE) cells. Cell fate patterns during this stage range from an alternating pattern at the beginning to the separation of Epi and PrE at the end. Several mechanisms guiding this decision and pattern formation have been proposed, including intra- and intercellular signalling, cell division and cell sorting. The current understanding is that signalling generates the cell fates and subsequent sorting introduces the spatial cell fate separation. We used agent-based modelling to investigate whether cell differentiation and cell sorting can act concurrently and how their relative contributions to pattern formation may change over time. Comparing our model to experimental data for mouse blastocysts and ICM organoids, we find two mechanistic regimes that can produce the experimentally observed spatial separation: (i) simultaneous long-range intercellular signalling and cell sorting, and (ii) a gradual transition from short-range signalling to cell sorting, in which the timing is mediated via reducing cell fate plasticity. While the second agrees better with existing experimental evidence for late blastocysts, the first might still be relevant for early and mid blastocysts. Together, our results refine the sequential view of Epi/PrE patterning by showing that fate specification and cell sorting can be dynamically coupled, with their relative contributions changing over the course of blastocyst development.

## 1. Introduction

Patterns are present at every level of biological organisation, from macroscopic markings on animal skin to microscopic arrangements of cells. Even at the subcellular level, ordered structures are prevalent, such as the lipid bilayer of the cell membrane or the base-pairing of DNA. Understanding the underlying mechanisms of pattern formation is therefore a central question in biology. In animals, many of these patterns emerge during early development.

One example at the cellular level is the second cell fate decision in mammalian preimplantation development. During this process, cells of the inner cell mass (ICM) differentiate into either epiblast (Epi) or primitive endoderm (PrE) cells and progressively segregate to form a spatially organised pattern [1]. Later, Epi cells give rise to the embryo, while PrE cells contribute to the yolk sac. This process has been well studied in early mouse embryos, where initially, all ICM cells coexpress the transcription factors NANOG and GATA6. Over time, this co-expression resolves as one factor becomes dominant and the other is downregulated, committing cells to either the Epi (NANOG expressing cells) or PrE (GATA6 expressing cells) fate. The spatial arrangement of the cell fates transitions from a random distribution through the formation of local clusters to the eventual separation of Epi and PrE cells [1], with a final PrE to Epi cell ratio of approximately 3:2 [2].

Several mechanisms have been proposed to contribute to the patterning during the PrE vs Epi decision [1], including intercellular signalling [3], cell division [4] and cell sorting [5, 6, 7, 8]. However, the relative importance and timings of the different mechanisms remain unclear. Because these processes occur simultaneously and influence each other, experimental analyses are complicated and mathematical modelling provides a valuable complementary strategy.

In our previous work, we investigated subsets of these mechanisms. We performed quantitative analyses of the three-dimensional cell fate patterns in mouse blastocysts and in ICM organoids [9, 10, 11, 12]. These organoids are three-dimensional aggregates derived from reprogrammed embryonic stem cells [13] that recapitulate key aspects of ICM patterning [9]. Combining the experimental data with mathematical modelling, we have shown that stochastic initial cell fate assignment followed by cell division with fate heredity is sufficient to reproduce local clustering, as observed in mid-stage ICM organoids [4]. In a separate approach, combining a gene regulatory network (GRN) with intercellular signalling that extends beyond direct neighbours reproduced both the local cell fate clusters and the correct PrE:Epi cell ratio in mouse embryos [14, 11]. However, neither model reproduces the engulfing pattern in late-stage ICM organoids or the clear segregation of cell fates observed in the late blastocyst. This suggests that additional mechanisms, or specific interactions between mechanisms, are required to reproduce the late stage pattern.

A candidate mechanism is cell sorting driven by differences in physical properties between cell types. The differential adhesion hypothesis [15] proposes that segregation arises from differences in effective surface tension between cell types, leading to energetically favourable configurations. Alternative proposals emphasise differences in mechanical properties such as elasticity [16] or cell surface fluctuations [7]. Importantly, these sorting models assume that the differences in physical properties are constant, resulting in different categories of cells.

Building on this categorical idea, previous approaches have shown that a sequence of first cell fate specification and then differential adhesion can generate the cell fate separation observed in late blastocysts [6, 5, 8]. A variation of differential adhesion relies on continuous expression levels of cell adhesion molecules (CAMs), allowing adhesion strengths to depend dynamically on gene expression regulated by an underlying GRN [17]. This formulation can be used to establish a direct mechanistic link between transcriptional regulation, in our case NANOG and GATA6, and physical cell sorting. Furthermore, it allows for cell fate specification and cell sorting to happen simultaneously in the model.

ICM cells remain uncommitted in early and mid blastocysts, and the timing of differentiation can be altered [18]. Furthermore, they exhibit a high degree of cell fate plasticity [19]. The cells can respond to environmental signals and adjust the intracellular concentrations of NANOG and GATA6 to switch between the two fates. As development progresses, the degree of cell fate plasticity decreases, and the cells irreversibly commit to Epi or PrE fate [20, 21].

We investigated how transcription regulation, intercellular signalling, cell division and adhesion-based sorting, linked to continuous NANOG and GATA6 expression levels, interact when operating simultaneously in a three-dimensional system. We explored how the degree of cell fate plasticity affects the final cell distribution pattern. Employing an agent-based centroid model, we systematically activated and deactivated the individual mechanisms and varied their relative strengths and temporal onsets. Analysing the resulting spatial patterns, we identified two mechanistic regimes that reproduce late-stage Epi/PrE spatial organisation: either (i) simultaneous long-range signalling and adhesion-based sorting, or (ii) a sequence of short-range signalling and cell sorting, induced by a reduced degree of cell fate plasticity.

## 2. Methods

We investigate how the interplay of transcriptional regulation, intercellular signalling, cell division, adhesion-based sorting, and cell fate plasticity affects cell differentiation patterns in ICM organoids and the ICM of mouse blastocysts. We built on our existing agent-based model, in which ICM organoids and the ICM are represented as three-dimensional aggregates of cells modelled as spheres with a centroid and a radius [14]. The model includes logistic growth dynamics for the radius and allows cells to divide, with the probability of division increasing with cell radius [11]. Upon division, the cellular volume is divided evenly between the daughter cells.

Cell fate depends on the concentrations of the two transcription factors *U* and *V* in each cell. Considering the Epi/PrE decision, *U* corresponds to NANOG and *V* to GATA6. The model incorporates a gene regulatory network (GRN) (Figure 1) for each cell with mutual inhibition and auto-activation of the two transcription factors. Cells communicate via a signal *S*, which is dependent on *U* . The signal activates *V* and inhibits *U* in neighbouring cells. This leads to the cells becoming either *U* - or *V* -positive during the simulations, depending on the received signal. In this model, autocrine signalling is not considered. Upon cell division, daughter cells inherit the same concentrations of *U* and *V* as the mother cell. Cellular movement is governed by a combination of Brownian motion and mechanical interactions between the cells, capturing both stochastic and deterministic aspects of cell dynamics.

**Figure 1.**
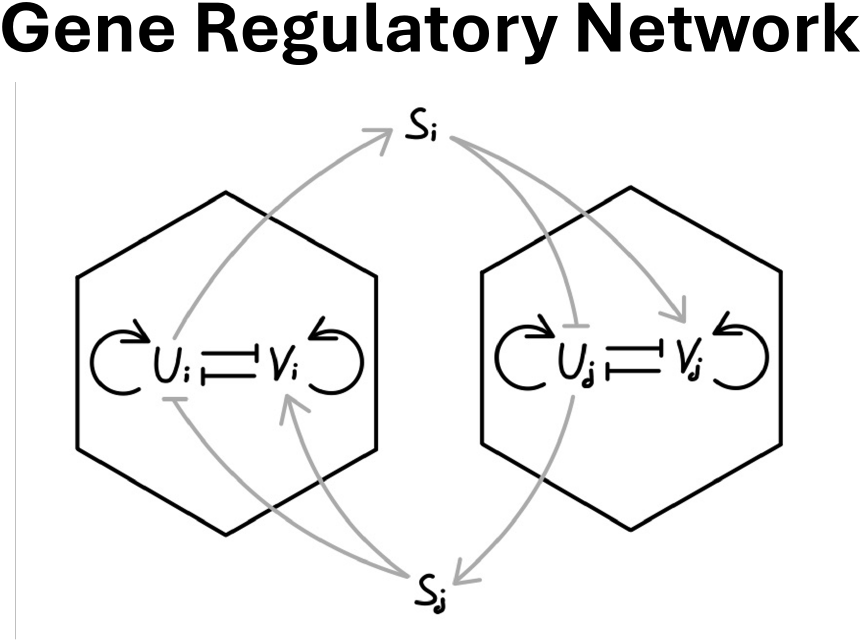
Illustration of GRN and cell-cell signalling for the two cells *i* and *j*, transcription factors *U* and *V* and signal *S*. Adapted from [14].

We simulate cell differentiation with different combinations of cell division, cell sorting and reduction in cell fate plasticity and discuss the emerging cell fate patterns in the context of our experimental data [9, 11] to investigate the influence of these factors on pattern formation.

### 2.1 Transcription

The transcription in the model is based on our previously developed system of ODEs [14]. The change of the two transcription factors *U* and *V* in cell *i* is given by:

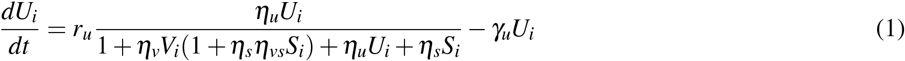

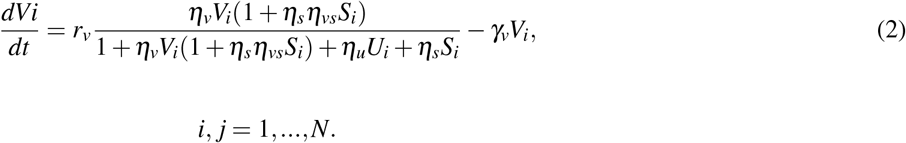

The change of concentrations over time is dependent on the degradation rates *γ*, the production rates *r* times the total binding probability as given by the fraction, where *η*_*x*_ denotes the statistical weights for the different binding events. At the single-cell level, the system has four steady states: the trivial *U*_*i*_ = *V*_*i*_ = 0 and depending on the signal level *S*, either the cells are *U* or *V* positive, whereby the other transcription factor is almost not expressed at all, or the levels of both are equal [14].

### 2.2 Cell fate plasticity

Cell fate decisions become irreversible at some point in development. In the mouse blastocyst, signalling has a decreasing influence on cell fate patterns over time [1]. We incorporate this aspect in Model D (see below) by limiting cell type plasticity. This is regulated by choosing a cut-off value *c*, such that the transcription level of *U*_*i*_ or *V*_*i*_ of a cell is set to zero if *U*_*i*_,*V*_*i*_ *< c*. At this point, a cell is fixed to be either *U* - or *V* -positive. In other words, if the concentration of one transcription factor inside the cells is too low, it is set to zero, stopping the cell from switching its cell type again. Changing *c* allows for fine-tuning the timing between signalling and transcription on the one side and sorting on the other. When *c* is reduced, cells take longer to reach the irreversible steady state, with either *U* or *V* concentration being high. This increases cell-fate plasticity, giving cells more time to respond to their local environment. If c is increased, cells have less time to respond to the local environment, and their fate is quickly locked. Since translating *c* into biology is difficult, we choose to test a wide range of values.

### 2.3 Signal

#### 2.3.1 Cell neighbourhood

We assume the signalling molecule is transmitted directly from cell to cell rather than freely diffusing through the aggregate. Hence, cells need to be in close contact with each other. To determine the cell neighbourhood, the aggregate is represented as a spatial cell graph [22]. Each cell is represented by a node, which has the spatial position of its centre of mass. We build the graph via a Delaunay triangulation, which connects each point to its neighbours in space, and then we eliminate all edges whose distance exceeds the sum of the radii of the two cells. The resulting graph consists of the centres of all cells as nodes, with edges indicating physical contact between cells in the model.

#### 2.3.2 Signal distribution

Cells can receive signals not just from their direct neighbours but also from cells further away. The signal strength depends on the distance *g*_*k, j*_ of two cells *k* and *j* on the cell graph and the level of *U* in the sending cell. The spatial decay of the signal is controlled by the parameter *q* ∈ (0, 1). Incorporating all this, *S* is defined as

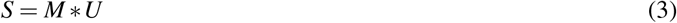

with

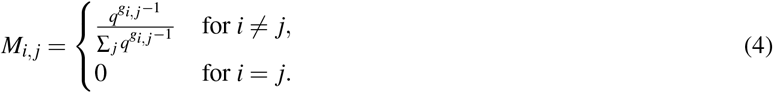

Hence, all emitted signal *S* is absorbed by other cells in the system. This signal scaling differs from our previous scaling [14], which led to an imbalance between the emitted and absorbed signals in the model.

### 2.4 Cell growth and cell division

As in our previous models, we assumed logistic cell growth and a stochastic, symmetric cell division along a randomly oriented axis ([11, 14]). The change in cell radius *r* is given by

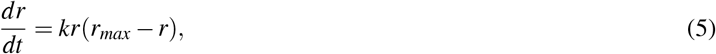

where *k* denotes a constant growth rate. Hence, for an initial cell radius *r*(*t*_0_) = *r*_0_, we get

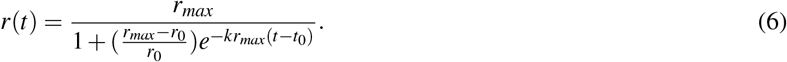

For cell division, we use the cumulative distribution function of the truncated normal distribution

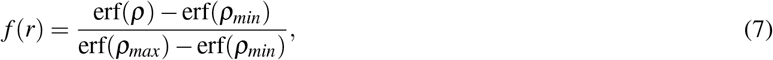

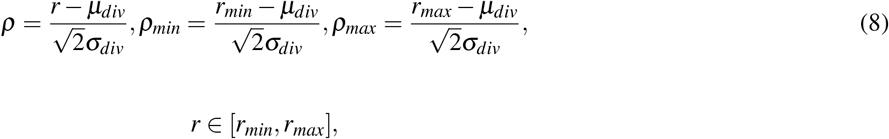

where *µ*_*div*_ and *σ*_*div*_ are the mean and standard deviation of this distribution. We chose 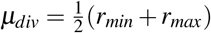. The function erf is the error function defined as

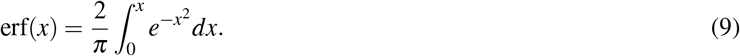

In addition, we set

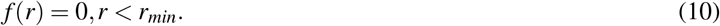

Hence, no cell divides up to a radius *r*_*min*_. Including previous attempts to divide, we obtain the division probability of cell *i* with radius 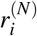 at time step *N* as

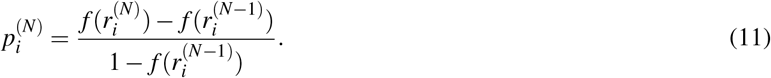

Following division, we use mass/volume conservation and assume symmetric cell division. Hence, the radii of the daughter cells *r*_1_ and *r*_2_ are given by

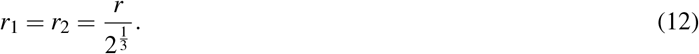

### 2.5 Cell movement and sorting

Cells in the model move by Brownian motion as well as due to intercellular physical interactions. The change of position *x*_*i*_ for cell *i* over time is then given by a combination of the forces *F*_*i, j*_ between the cell *i* and its neighbouring cells *j* and random movement *ε*_*i*_ drawn from a normal distribution:

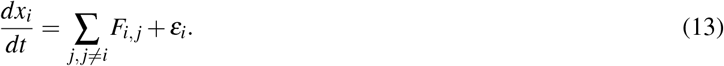

In the model, if two cells come in contact, they exert forces on each other. For short distances, these forces are repulsive; for longer distances, they are attractive. Above a given distance, the cells no longer exert any forces on each other. To incorporate differential adhesion based on continuous expression of the transcription factors inside the cells, we used an adapted version of the Morse potential [23], where *F*_*i, j*_ depends on the Euclidean distance *d*_*i, j*_ between two cells and their radii *r*_*i*_ and *r* _*j*_ such that

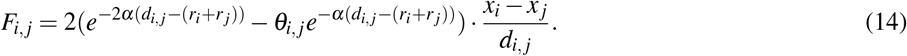

The constant *α* depicts the stiffness of the Morse potential. The adhesion *θ*_*i, j*_ is different for each cell pair and depends on their level of *U* and *V* such that

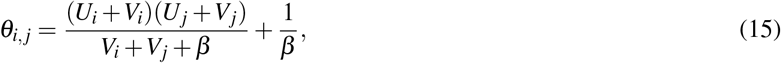

with a scaling parameter *β* > 0. The extreme cases of *θ*_*i, j*_ are illustrated in the following, while resulting forces are shown in the appendix (Figure S1):

| $U_i$ | $V_i$ | $U_j$ | $V_j$ | $\theta_{i,j}$ |
| --- | --- | --- | --- | --- |
| 1 | 0 | 1 | 0 | $\frac{2}{\beta}$ |
| 1 | 0 | 0 | 1 | $\frac{1}{1+\beta} + \frac{1}{\beta}$ |
| 0 | 1 | 0 | 1 | $\frac{1}{2+\beta} + \frac{1}{\beta}$ |

Hence, the order of affinity is *UU* > *UV* > *VV* , which has been shown to promote engulfing patterns [17] and has previously been used in a model for cell sorting in the mouse embryo [5]. By setting *θ*_*i, j*_ = 1 in (14), we obtain the basic form of the Morse potential, where no sorting happens but cells still move due to cell division, growth and Brownian motion.

### 2.6 Simulation

We conducted simulations with different combinations of mechanisms, starting with just signalling and transcription and building up complexity (Table 1).

**Table 1.** Model setups used in this study.

| Model label | signalling<br>+ transcription | cell division<br>+ movement ($\theta_{i,j} = 1$) | cell sorting<br>($\theta_{i,j}$ from (15)) | reducing cell<br>plasticity |
| --- | --- | --- | --- | --- |
| Model A | X |  |  |  |
| Model B | X | X |  |  |
| Model C | X | X | X |  |
| Model C2 |  | X | X |  |
| Model D | X | X | X | X |

In Model A without cell division, simulations were conducted on a fixed 300-cell 3D aggregate. The aggregate was initiated with a single cell and allowed to grow to its final size before transcription commenced. Unless otherwise stated, initial values for *U* and *V* are given by

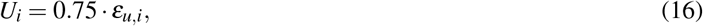

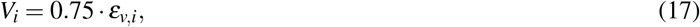

where *ε*_*u,i*_ and *ε*_*v,i*_ are drawn from a normal distribution with mean zero and standard deviation 0.1. Aggregates were simulated until the transcription in the cells reached a steady state. The aggregate was considered to be in a steady state if

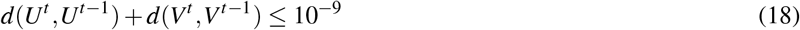

where *d*(., .) denotes the Euclidean distance. The superscripts *t* and *t* − 1 refer to two consecutive time points of transcription factors *U* and *V* of all cells.

Simulations with cell division (Models B, C, C2 and D) start with five cells in proximity to each other with the same conditions for the initial expression level as before. The simulations end when the aggregate reaches a size of 300 cells, regardless of whether the transcription is in steady-state or not, to be comparable in size to the simulations for Model A.

Model C2 serves as a control for the influence of cell sorting, so no transcription or signalling takes place. Here, upon initialisation, *U* and *V* positive cells were set up in a ratio comparable to the final numbers of *U* and *V* positive cells in the other model versions, and cell type is inherited during cell division.

The remaining default parameter values are summarised in Table 2. For parameter scans, we performed 50 replications for each combination of parameter values. Parameter values were chosen such that all model variants exhibit comparable final cell fate ratios U+:V+ of 2:3, matching the experimentally measured ratio of Epi:PrE of 2:3 (Figure S2, [2]).

**Table 2.** List of model parameters, including values and description. Parameters were adapted to allow a steady state concentration of 1 for *U* or *V* . Adapted from [11].

| parameter | value | description | equation |
| --- | --- | --- | --- |
| $\gamma_U$ | 1 | Decay rate of $U$ | (1) |
| $\gamma_V$ | 1 | Decay rate of $V$ | (2) |
| $r_U$ | 1 | Transcription rate of $U$ | (1) |
| $r_V$ | 1 | Transcription rate of $V$ | (2) |
| $-\Delta\epsilon_S$ | 0.849 | Energy difference w.r.t. binding of $S$ | (1),(2) |
| $-\Delta\epsilon_V$ | -7.10 | Energy difference w.r.t. binding of $V$ | (1),(2) |
| $-\Delta\epsilon_{VS}$ | -0.849 | Energy difference w.r.t. binding of $V$ and $S$ | (1),(2) |
| $-\Delta\epsilon_U$ | -6 | Energy difference w.r.t. binding of $U$ | (1),(2) |
| $q$ | $\in (0, 1)$ | Dispersion parameter for distance-based signal | (4) |
| $r_{max}$ | 1 | Maximum cell radius | (5),(7),(8) |
| $r_{min}$ | 0.9 | Minimum cell radius for division | (8),(10) |
| $k$ | 0.1 | growth rate of the cell radius | (5) |
| $\alpha$ | 2 | cell stiffness | (13) |
| $\sigma$ | 0.7 | optimal distance between two cells in contact as a fraction of the sum of their radii | (13) |
| $\beta$ | 1 | scaling constant for the mutual affinity | (14) |
| $F_0$ | 0.1 | scaling of Morse potential | from [11] |
| $c$ | $10^{-7}$ | transcription cut off | section 2.1 |

### 2.7 Metrics

We use Moran’s index, a measure of spatial autocorrelation, to quantify the resulting cell fate patterns [24]. The Moran’s index *I* for *n* cells is given by:

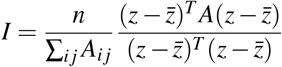

where *z* represents the cell fate with *z*_*i*_ = 1 if *U*_*i*_ > *V*_*i*_ else it is 0. The mean is given by 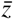, and *A*_*i j*_ is the adjacency matrix of the cell graph that represents the cell neighbourhood.

The range for the Moran’s indices for ICM organoids was obtained as described in [12].

For the range of Moran’s indices for the ICMs of late blastocysts, we used the values for the NANONG patterns from [11]. The values for the GATA6 patterns are comparable. They are between −0.1 and 0.7.

### 2.8 Implementation

Simulations and analyses were performed in Julia [25] and its package Dataframes.jl. All visualisations have been created using the package Plots.jl [26]. The code for the simulations and for creating all the figures is available on Github.

## 3. Results

### 3.1 Signalling can induce spatial cell fate separation

The aim of our study is to investigate the interplay of intercellular signalling, cell division, cell sorting and cell fate plasticity during Epi vs PrE differentiation. To this end, we simulated different combinations of the four mechanisms and analysed the resulting cell fate patterns. We started with Model A, which incorporates transcription of two transcription factors *U* and *V* and intercellular signalling on an aggregate with fixed cell number and positions (Table 1). For the intercellular signalling, the signal spreads to cells beyond the nearest neighbours, depending on a dispersion parameter *q* ∈ (0, 1). Increasing *q* increases the range of signal dispersion. For a value of *q* close to zero, the signal reaches only the direct neighbours on the cell graph. For *q* close to one, the signal is equally distributed among all cells in the system, except for the sender, regardless of distance. We examined the patterns arising from Model A. We used a size- and position-fixed aggregate of 300 cells with high concentrations of *U* and *V* in each cell as an initial condition. Simulating our model until steady state and observing the spatial distribution of *U* - and *V* -positive cells shows that our results are comparable to those from our previous model [14] (Figure 2), despite adapting the signal scaling (see Materials and Methods for details). A small dispersion parameter *q* leads to an alternating pattern of *U* -and *V* -positive cells, while there is an engulfing pattern for large *q* values. For intermediate values, we observe local clustering. It is possible to get an engulfing pattern only with transcription and signalling, but compared to the late stage ICM organoids, the positions of *V* -positive (Epi) and *U* -positive (PrE) cells are inverted.

**Figure 2.**
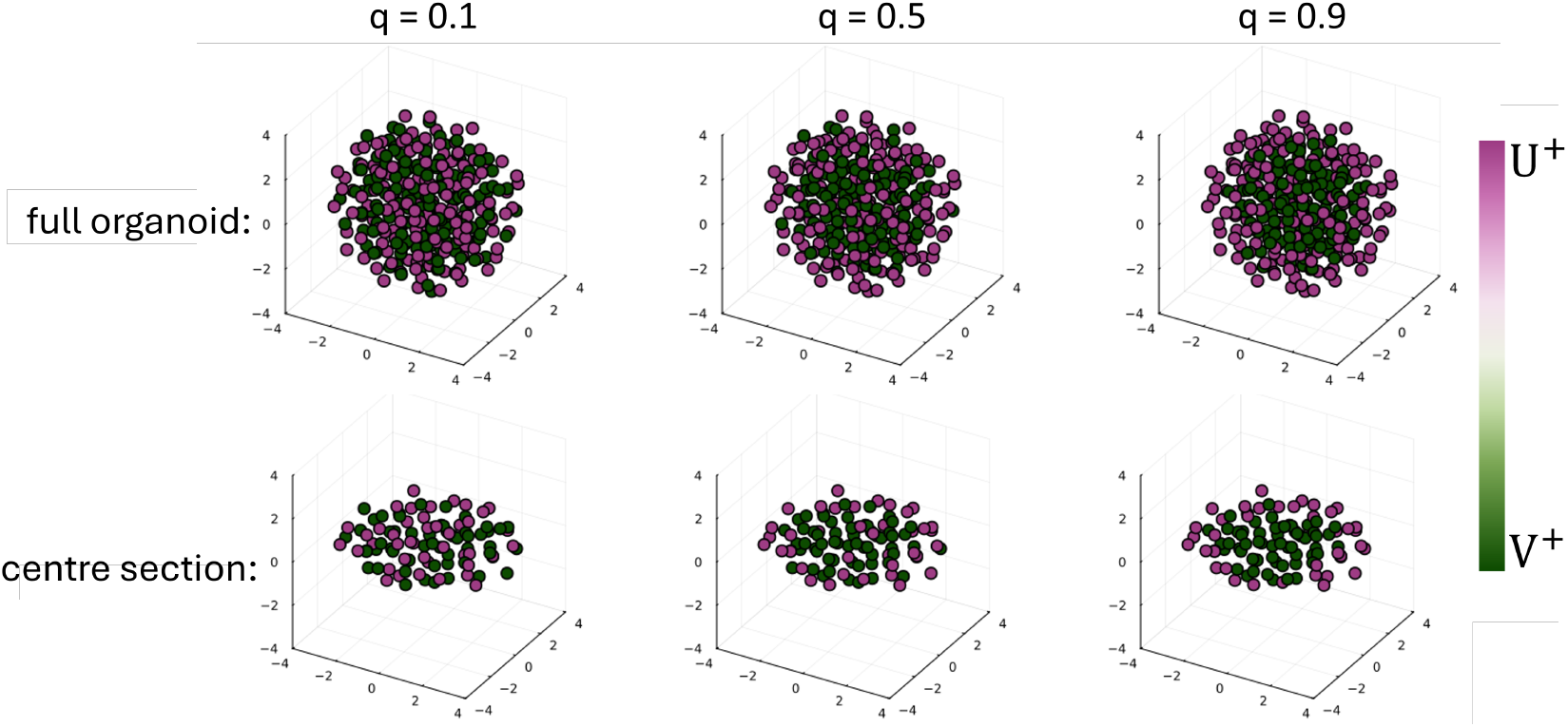
Patterns arising from Model A (signalling). Example simulation results for three different values of the dispersion parameter *q* in Model A and the remaining parameter values from Table 2. Each simulation is performed on the same 300-cell aggregate with high but randomised initial conditions for each cell’s *U* and *V* levels. Nodes of the cell graph, representing the centres of the cells, are plotted. Nodes are coloured according to the steady-state transcription level of *U* . The complete aggregate is displayed in the top row, while the lower row shows the corresponding cross-section through the centre.

For a quantitative assessment of the three-dimensional cell fate patterns, we employed the spatial autocorrelation coefficient Moran’s index *I* [24, 11, 12]. It takes values between −1 and 1. Highly negative values indicate an alternating pattern of two cell types, values close to 1 indicate spatial separation, while values close to zero describe random patterns. We use this measurement to systematically investigate the influence of the signalling range *q* on the resulting patterns of Model A (Figure 3). We observe an increase in Moran’s index from −0.2 for *q* = 0.1 to 0.4 for *q* = 0.9. Most of the increase in Moran’s *I*, which indicates increasing separation of the *U* and *V* positive cells, happens until *q* = 0.7. Increasing *q* further only slightly increases Moran’s *I*. For each tested parameter, the variance is low, indicating the resulting patterns are robust.

**Figure 3.**
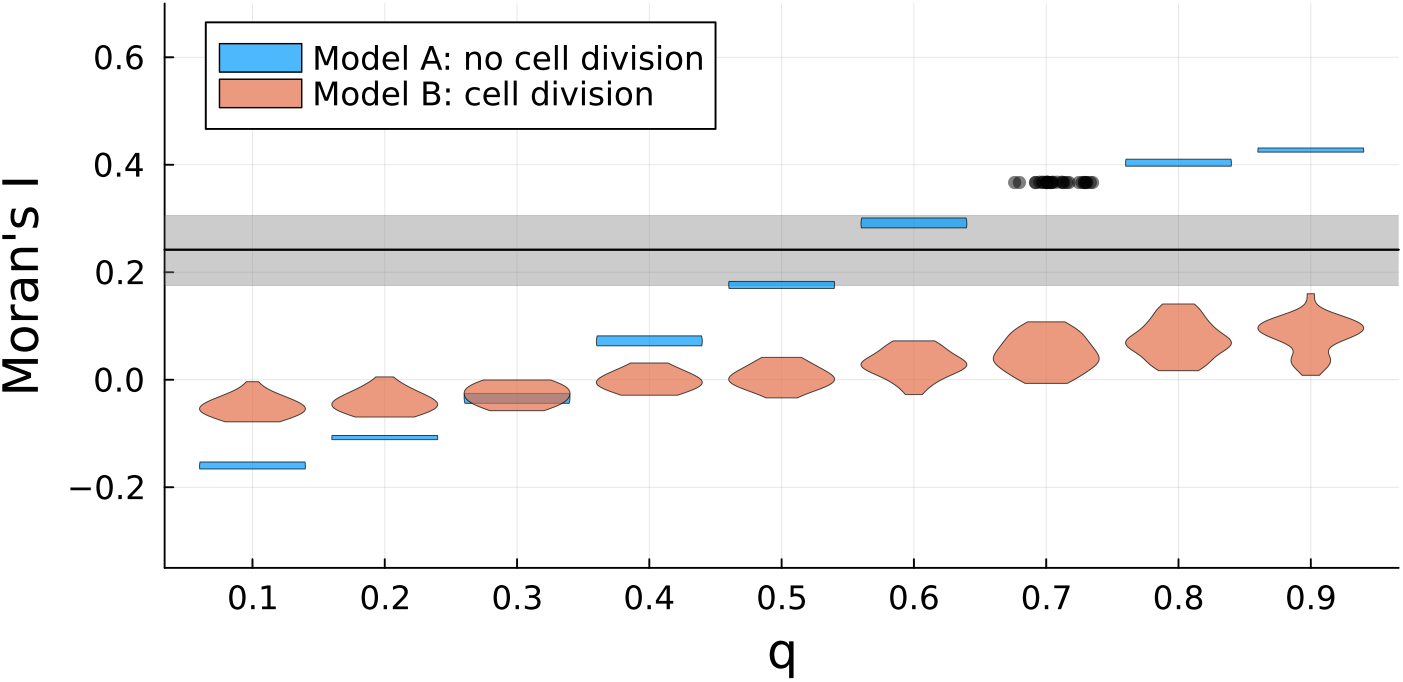
Influence of signalling range on spatial cell fate separation for Models A (signalling) and B (signalling+cell division). Violin plots of the Moran’s index against the dispersion parameter *q* for Models A and B. Each violin consists of 50 simulations. For Model A and *q* = 0.7, individual simulations are visualised as dots, since the variance in Moran’s *I* is too small for a violin. The remaining parameter values are summarised in Table 2. The grey line shows the median experimental value for late ICM organoids [12]. The grey area marks the interquartile range, which represents the middle 50% of the data.

To put the simulation results into context, we compared them to our experimental measures for late ICM organoids [12] (Figure 3), which have a median Moran’s index of 0.24 with a first quartile Q1 of 0.18 and a third quartile of 0.31. The range is 0.009 to 0.41. The ICM of late blastocysts does not exhibit an engulfing pattern, but Epi and PrE are adjacent. However, the Moran’s indices are comparable with a median of 0.22, Q1 of 0.06, Q3 of 0.37 and a range of −0.17 and 0.62 [11]. We observe that for intermediate *q*, the simulation results overlap with the central 50% of the distribution of the experimental data, but do not reach the largest value of the mouse blastocysts. The agreement with respect to Moran’s index is not in contrast to the discrepancy in ordering of cell fates (Figure 2, because the Moran’s index does not take this order into account.

As a next step, we included cell division in the model, which also requires allowing cell movement based on Brownian motion and mechanical interactions to maintain the coherence of the simulated aggregates (Model B). Now, simulations start with a five-cell aggregate and terminate once the aggregate reaches 300 cells, regardless of the state of other variables. Since all mechanisms are active simultaneously in the simulation, signalling and transcription begin at the start of the simulation. We calculate the Moran’s index for the aggregate at the final time point and find that for all tested dispersion parameters, it is close to zero, indicating less cell fate separation than for Model A (Figure 3). There is still an increase in Moran’s *I* as *q* increases, but it is much smaller and more linear than in Model A. Therefore, cell division seems to dominate over intercellular signalling in regard to pattern formation. Also, resulting patterns are not as conserved in Model B as before, since the variance in Moran’s *I* for each parameter is higher than in Model A. This effect is most likely due to cell division and the resulting cell movement, which introduces noise into the spatial positioning of cells. Finally, the simulation results do not overlap with the interquartile range of the experimental values for late ICM organoids. In summary, signalling and transcription can generate varied patterns, ranging from alternating to engulfing and with Moran’s indices comparable to the mouse ICM. However, the resulting model patterns are in reverse order compared to the late stage experimental data. Including cell division weakens cellular separation.

### 3.2 Sorting can induce experimentally observed engulfing pattern

So far, we showed that signalling can generate different cell fate patterns as indicated by the distribution of the Moran’s indices, but also that cell division reduces this effect. Since cell sorting is a key aspect during development as cell fate decisions are made and cells locate in their final location, we include it as a third process in Model C (Table 1).

To study cell sorting in the context of movement and division, we extended our model based on an affinity-based ordering of cell–cell interactions [5], which suggests the order of adhesion strength: *UU* > *UV* > *VV* . Based on this, we developed a quantitative scheme for differential adhesion that depends on the *U* and *V* levels within cells (see Materials and Methods for details). We started the simulations with five cells and terminated once 300 cells were reached, regardless of the other variables. The final cell fate patterns for values of the dispersion parameter *q* between 0.1 and 0.9 were again quantified employing Moran’s *I* (Figure 4).

**Figure 4.**
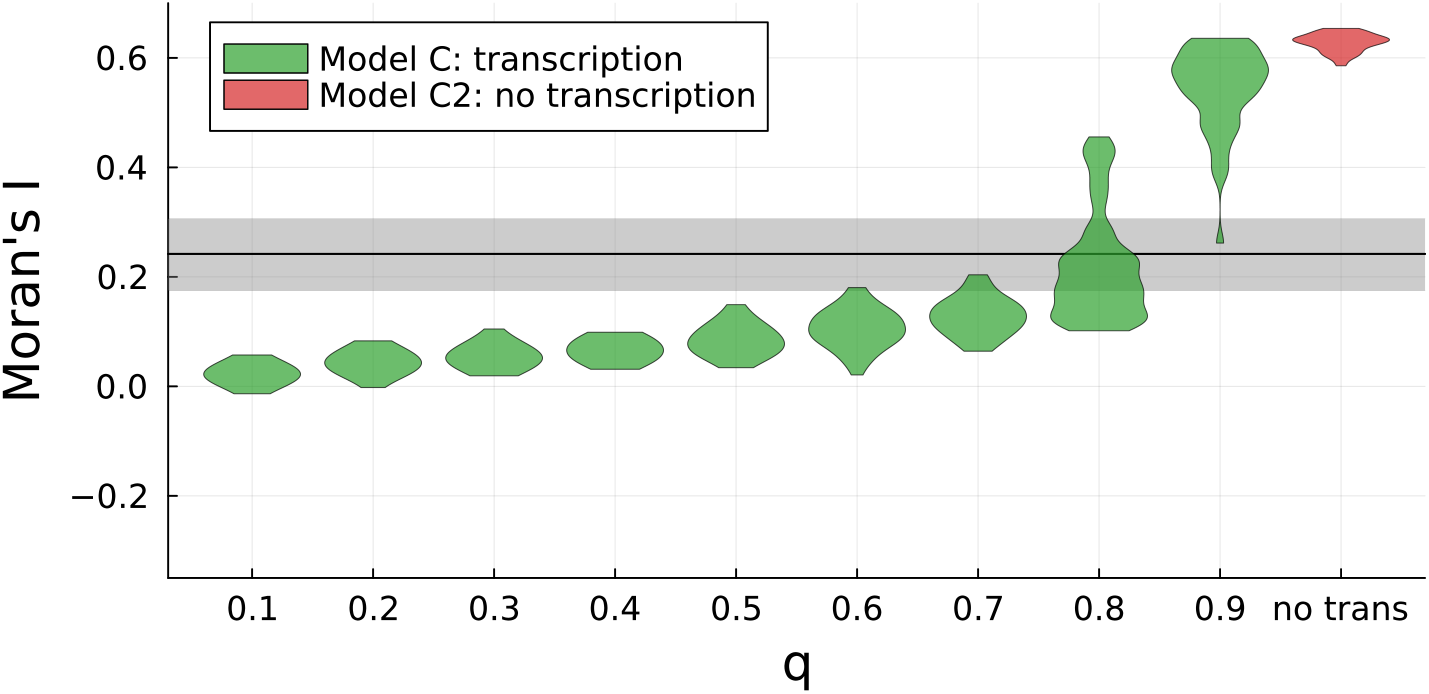
Influence of signalling range on spatial cell fate separation for Models C (signalling+division+sorting) and C2 (control without transcription and signalling). Violin plot of the Moran’s index for simulations of Models C and C2. Each violin consists of 50 simulations. The dispersion parameter *q* was changed from 0.1 to 0.9. Other parameter values are given in Table 2. The grey line shows the median experimental value for late ICM organoids [12]. The grey area marks the interquartile range, which represents the middle 50% of the data.

We find that for *q* ≤ 0.7, the results of Models C and B are comparable, with overall slightly higher Moran’s *I* for Model C, indicating that cell sorting only has a small effect on cell fate patterning here. For *q* ≥ 0.8, Moran’s index increases strongly with a big jump from *q* = 0.8 to 0.9, indicating a big increase in separation of the two cell fates. There, we further observe a much larger variance than for Model B. For *q* = 0.9, Moran’s index assumes its largest value for Model C, which is 0.6 and even higher than the 0.4 in the static Model A, showing that the combination of sorting and signalling can lead to stronger cell fate separation than signalling alone.

A comparison of the modelling results with Moran’s indices for ICM organoids [12] shows that to obtain simulation Moran’s indices within the interquartile range of the experimental data, larger *q* values are required than for Model A. Furthermore, for *q* = 0.9 the largest Moran’s index of 0.6 of the late mouse blastocysts can be obtained with Model C.

To investigate the effect of sorting without signalling and transcription, we simulated Model C2. The results show a similar Moran’s *I* as Model C for *q* = 0.9, but a significantly smaller variance (Figure 4). This indicates that pattern formation is less stable in the model when signalling and sorting are combined, compared with the patterns resulting from the individual processes.

To investigate whether the organisation of the resulting separated pattern is as expected with *U* positive cells (Epi) on the inside, engulfed by *V* positive ones (PrE), we plotted the spatial distribution of the *U* and *V* levels in simulated aggregates for Model C for *q* = 0.1, 0.5 and 0.9. Aggregates tend to be smaller in diameter than in simulations without sorting, since adhesive forces are higher than in Model B. For *q* = 0.1 and 0.5, there are cells shown in an intermediate state, where they express both transcription factors at an intermediate level (indicated by their white colour (Figure 5)). For the high *q* value simulation, the aggregate shows the expected pattern of *U* -positive cells (Epi) engulfed by *V* -positive ones (PrE), which is the reversed pattern compared to Model A (Figure 2). In summary, cell sorting based on the implemented affinity order in combination with long-range signalling yields Moran’s indices comparable to experimental measures, and the spatial arrangement of the two cell types is in the expected order.

**Figure 5.**
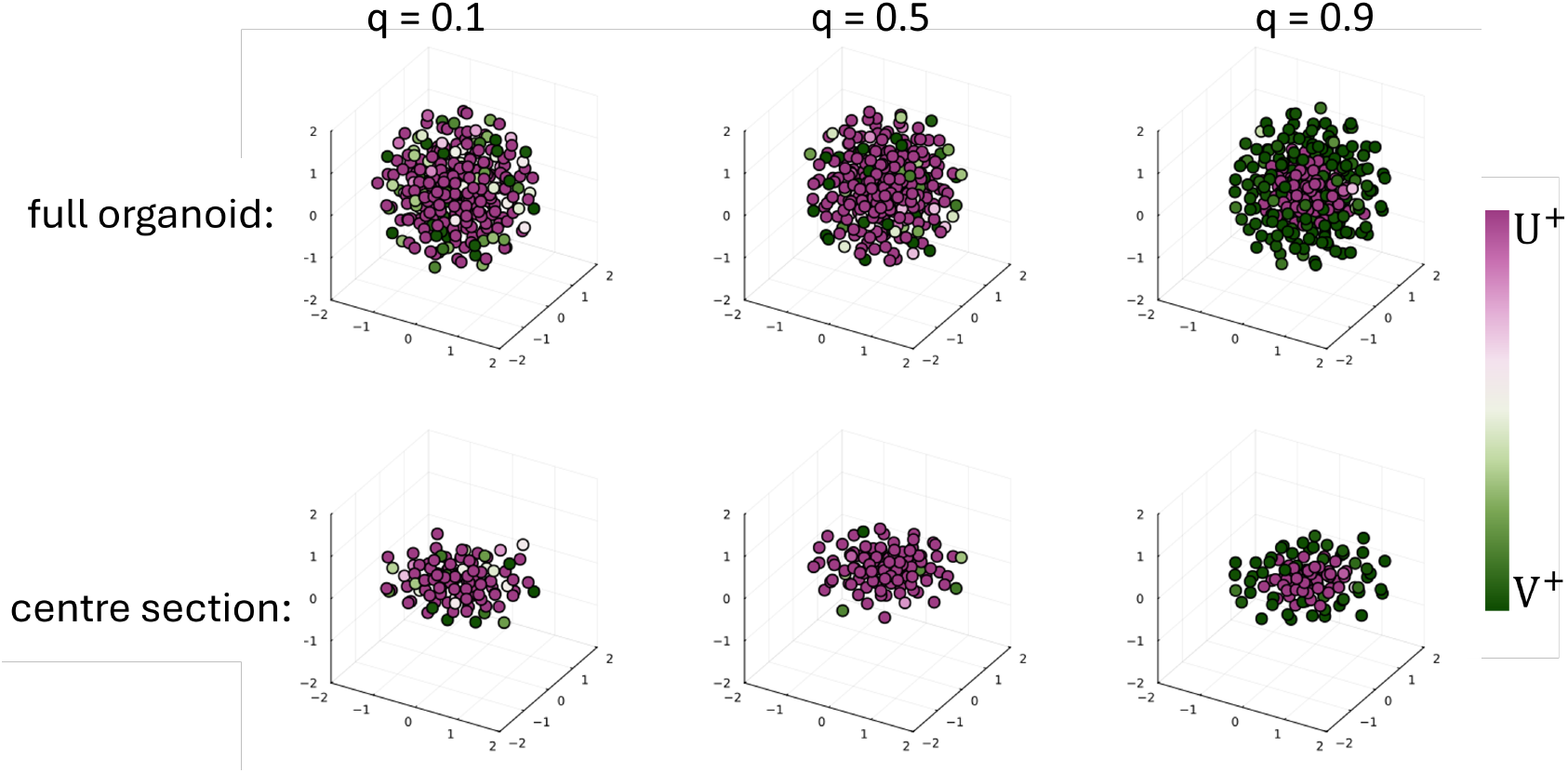
Patterns arising from Model C (signalling+division+sorting). Example simulation results for three different values of the dispersion parameter q in Model C and the remaining parameter values from Table 2. Simulations start with five cells, with high expression levels of *U* and *V* . Each aggregate grows until it reaches a size of 300 cells. Plotted are just the nodes of the cell graph, representing the centres of cells. Nodes are coloured depending on the transcription level of *U* . The complete aggregate is displayed in the top row, while the lower row shows the corresponding cross-section through the centre.

There is still the question of why sorting does not achieve this kind of cell fate separation for the other parameter values of *q*, i.e. shorter-range signalling. One possible explanation is that signalling and sorting might compensate for each other when sorting has only a small effect on the separation of cell types. Since cells are not committed, they can change their fate when placed in a new local environment due to sorting (Supplementary Figure S3). For Model C and low *q*, we observe the largest number of cell fate switches. In this scenario, there seems to be an endless cycle of sorting and cell fate switching.

### 3.3 Different mechanistic regimes form engulfing patterns

We next asked: What would be needed for a clear engulfing pattern for signal dispersion parameter values below 0.9? In vivo, signalling in the mouse blastocyst exerts a diminishing influence on fate patterns over time, and cell fate decisions become irreversible as development proceeds [27]. To add this restriction on cell fate plasticity to our model, we introduced a cut-off parameter *c* (Model D). As soon as the concentration of *U* or *V* falls below *c*, the respective value is set to zero. This then prevents the cell from producing any more of the corresponding transcription factor, effectively “locking” it into the other fate, regardless of the signal it receives from its neighbours. By changing *c*, it is possible to regulate cells’ plasticity, adjusting the time they have to make their final cell fate decision. If *c* is small, cells have higher plasticity. Hence, it takes longer for them to reach such a small concentration of *U* or *V* that they are fixed in the other fate. In other words, by changing *c*, we can adjust the time that cells remain uncommitted.

We used Moran’s *I* to quantify the resulting simulation patterns for different combinations of *q* and *c* in Model D (Figure 6 (a)). For the cut-off values *c* that we considered, simulated aggregates tend to have larger mean Moran’s *I*, indicating a higher separation of the cell types (Figure 6 **(a)**) than without including the reduction in plasticity (Model C, Figure 4). However, there is a clear effect from the choice of cut-off value *c* and dispersion parameter *q*.

**Figure 6.**
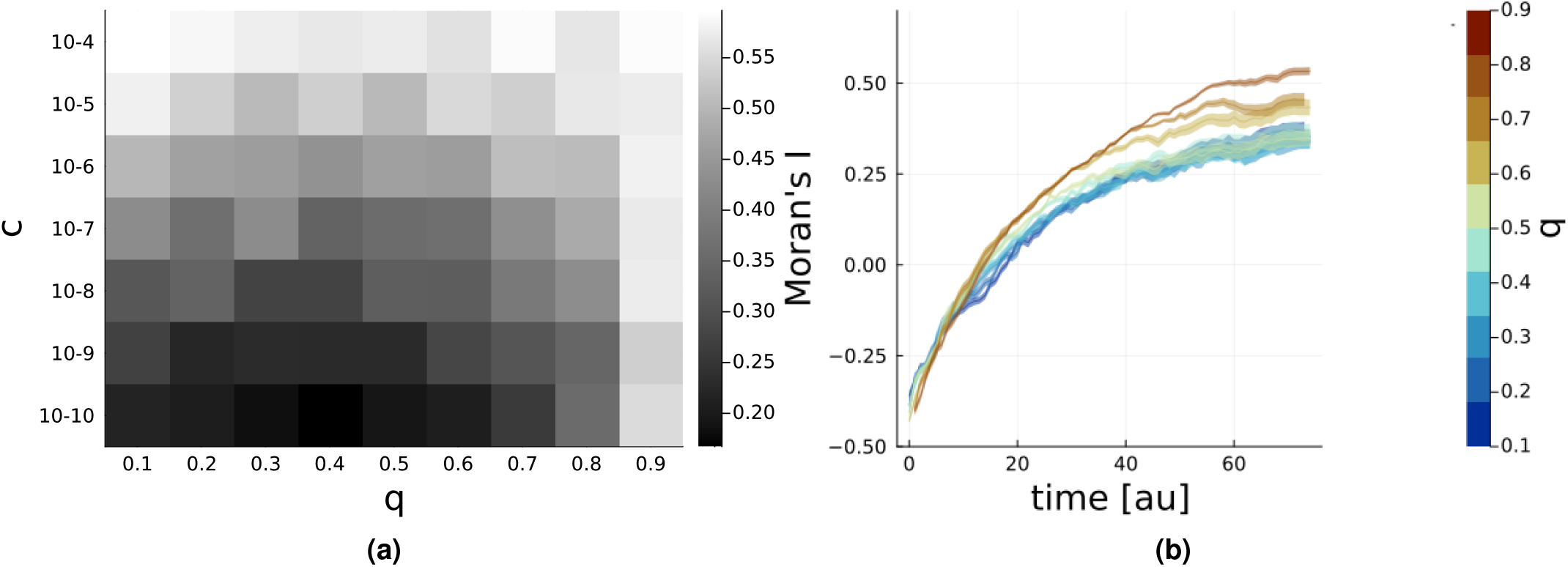
Influence of signalling range, cell fate plasticity and time on spatial cell fate separation for Model D (signalling+division+sorting+reduced plasticity) **(a)** Heatmap of the mean Moran’s index for Model D and combinations of the cut-off *c* and the dispersion parameter *q*. For each combination, 50 runs were simulated until the aggregate reached 300 cells. **(b)** Mean Moran’s *I* with standard error of the mean for Model D over time shown for different values of the dispersion parameter *q* and a fixed cut-off *c* = 10^−7^, while other parameter values are given in Table 2. For each *q* value, 50 simulations were run, starting with 5 cells and until the aggregate reached 300 cells.

For *q* = 0.9, cell fate separation seems to be mostly independent of *c*, since the mean of Moran’s *I* is always high (above 0.5). For large and intermediate *c* values, signal dispersion is not important for pattern formation, because Moran’s *I* is high in all these cases. When *c* is decreased, cells remain uncommitted longer and have more time to decide. Then, Moran’s *I* decreases and depends on signal dispersion *q*. For low *c* and *q* = 0.3 as well as 0.4, we obtain the lowest Moran’s indices of and an overlap with the interquartile range of the experimentally observed Moran’s indices for late ICM organoids [12]. Furthermore, all Moran’s indices obtained with Model D are within the upper half of the range of the experimental values for late ICM organoids and ICMs of late blastocysts. It is important to note that compared to Model C, the patterns obtained with Model D have a much higher variance for most *q* (Figure 4 and Supplementary Figure S4). In summary, for low cell type plasticity (large *c*) or large signal dispersion *q*, we get the highest Moran’s indices and hence the most separation of cell types. So far, we have focused on the patterns at the end of the simulations. Our modelling approach further enables the analysis of the dynamics of pattern formation throughout the time frame of aggregate development (Figure 6 (b)). For Model D and *c* = 10^−7^, initially, the mean Moran’s *I* is negative for all *q* values and increases over time. This trend is similar to that observed in the experimental results [12, 11].

To summarise, signalling on its own can induce cell fate separation but the cellular arrangement does not agree with experimental observations in mouse blastocysts and ICM organoids. Introducing sorting preferences, where *UU* interactions are favoured over *UV* and *VV* , results in the observed order of the engulfing pattern for a high signal dispersion. Limiting cell fate plasticity enhanced the separation of cell types at lower signal dispersion values.

## 4. Discussion

We studied combinations of multiple mechanisms to investigate how the spatial separation of Epi and PrE cells in late ICM organoids and late mouse blastocysts could arise. We expanded our previous ODE-based organoid model, which included signal dispersion and cell division [14, 11], to incorporate sorting via differential adhesion that depends on continuous transcription factor levels within cells. By additionally exploring the impact of modulating cell plasticity, we identified two possible mechanistic regimes: either long-range signalling and adhesion-based sorting act simultaneously or short-range signalling is followed by cell sorting. In the latter case, the timing is determined by the degree of cell fate plasticity.

We first consider the option of long-range signalling and adhesion-based sorting acting simultaneously. The GRN underlying transcription and signalling in our model has been used by us and others previously [14, 28]. It relies on auto-activation of NANOG and GATA6, mutual intracellular inhibition of the two, and a paracrine signal that inhibits NANOG and activates GATA6. For long-range signalling, this system can generate spatial cell fate separation to a degree similar to that in ICM organoids and mouse blastocysts. However, the order of arrangement of the cell is inverted with *V* -positive cells (Epi) surrounding *U* -positive cells (PrE). Introducing cell sorting in the model reverses the cellular arrangement to the experimentally observed *U* -positive cells (PrE) surrounding *V* -positive cells (Epi). Sorting in our modelling approach is based on the differential adhesion hypothesis. The concentration of adhesion molecules is directly dependent on the concentrations of the transcription factors *U* and *V* inside the cells. This is different from previous modelling approaches for Epi and PrE differentiation, which typically first assume a definition of the cell types Epi or PrE and then discrete values for the adhesion strengths [5, 6, 8]. However, our idea aligns with previous experimental work arguing that dynamic physical parameters are required to explain Epi and PrE sorting [7]. As candidates for driving differential adhesion, EphA4 and EphrinB2 have been proposed as a ligand/receptor pair [8]. Furthermore, P- and K-cadherins have been proposed to play a role, while E-cadherin does not seem to be involved [29, 30]. In addition to differential adhesion, several further mechanisms have been considered important in Epi/PrE patterning, such as cortical tension [29], cell surface fluctuations [7] and cell polarisation [31]. We expect sorting in the mouse blastocyst to involve multiple mechanisms. Identifying their relative importance could be an interesting aim for future modelling studies. This will probably require replacing the centroid-based approach with a vertex model to incorporate cell shapes and thereby more refined mechanical interactions [32]. The open question in this scenario is what the long-range signalling would be. The signalling pathway usually considered in this GRN is FGF/MAPK signalling. Previous work has shown that this pathway acts over a short range [3]. Hence, for this scenario to take place in the mouse blastocyst, either another signalling pathway has to be involved or the long-range FGF signalling could be modulated via FGF diffusion at the extracellular matrix, FGFR trafficking, fgf4 mRNA decay, cytonemes, microvesicles or cytoplasmic bridges, processes that remain to be investigated in this context [33, 34, 35]. If we combine adhesion-based cell sorting with short-range signalling, we do not get a clear separation of cell fates. In this setting, the model predicts that cells can switch types if placed in new microenvironments via sorting, potentially creating competing dynamics between adhesion-driven sorting and signalling-induced changes in differentiation. This aligns partially with experimentally observed GATA6 and NANOG trajectories that switch during a specific window in the Epi/PrE cell fate decision in mouse blastocysts [36].

We now turn to the second option: a sequence of short-range signalling and cell sorting, in which the timing is introduced by modulating cell fate plasticity. Assuming that the intercellular signalling in our GRN relies on FGF/MAPK, a short-range signal fits with previous results [3]. The sequential approach is also in line with previous models for Epi and PrE differentiation [5, 6, 8]. The main difference is that existing models typically assume a fixed time point at which the cell fates are decided and sorting begins. Such an abrupt switch is difficult to imagine in a biological system. We introduce a concept for implementing the sequential approach with a more gradual switch. In our scenario, not the whole system but rather individual cells switch from differentiation to fixed cell fate and the switching depends on their levels of *V* and *U* , i.e., NANOG and GATA6. We define this concept as loss in cell plasticity, where cell plasticity is the relative ease with which a cell can switch between potential fates in response to intra- and intercellular signalling. We took into consideration that, as development progresses toward the late blastocyst stage, the ICM cells undergo a progressive loss of plasticity, transitioning from a pool of flexible, bipotential progenitors to irreversibly committed Epi and PrE cells [19, 20, 2, 21]. In vivo, this reduction in plasticity is mirrored by a declining sensitivity to FGF/MAPK signalling; while early uncommitted progenitors can be fully redirected by modulating this pathway, specified cells eventually become insensitive to these external cues, effectively “locking in” their identity as the window of responsiveness closes by late blastocyst stage [20, 2]. The exact mechanism by which cell plasticity is regulated remains to be fully characterised. Our recent work on the activity of Wnt/*β* -catenin signalling in this context provides some evidence [18]. We showed that increasing *β* -catenin levels led to faster PrE differentiation, whereas in its absence, ICM cells remained uncommitted for longer. Hence, it might be possible that Wnt/*β* -catenin signalling is involved in the regulation of cell plasticity. Consistent with this, we found that high levels of cell fate plasticity (low values of *c*) in Model D yield weaker cell fate separation as observed in *β* -catenin mutant ICM organoids. This suggests that Wnt signalling could be involved in the timely exit from a poised state [18]. Focusing on the molecular cross-talk between Wnt/*β* -catenin and FGF/MAPK signalling would be interesting, as their cooperation might dictate the speed of the transition from bipotentiality to terminal commitment.

Considering the experimental evidence, the second scenario of a gradual transition of sequential cell fate decision to cell sorting is more likely for late-stage cell fate decisions. For the other blastocyst stages, we previously determined Moran’s indices with a median of −0.02, Q1 of −0.07 and Q3 of 0.03 for early blastocysts and a median of −0.02, Q1 of −0.07 and Q3 of 0.05 for mid blastocysts [11]. This is not compatible with the steep increase in Moran’s index over time observed for the sequence of short-range signalling and sorting (Figure 6). Taking into consideration how cell fate acquisition progresses, there could also be a combination of both mechanisms operating during blastocyst development. Our results are compatible with cells following long-range signalling and sorting at early stages, when there are fewer cells and sequential short-range signalling and sorting as development progresses. Furthermore, as cells acquire their final fate asynchronously [2], it is possible that within the embryo, both models can act simultaneously, depending on the cell.

For future comparisons between the model and experimental data, especially to investigate the dynamics of the system, it would be of interest to have time-series data on ICM organoids, with individual cells tracked in position and in NANOG/GATA6 expression levels over time. To our knowledge, there is currently no such data available. For mouse embryos, live imaging of transcription factors during the EPI/PrE cell fate decision in blastocysts has yielded interesting data [36]. Adaptation of this technique to ICM organoids would be of interest. Alternatively, adapting the model to include the trophectoderm and the blastocoel would allow more direct comparisons between the available experimental data and the modelling results.

So far, the Epi/PrE decision has been modelled as a sequential process of cell fate specification followed by cell sorting, with an abrupt switch between these two modes of action. Our approach refines this view by introducing a gradual transition between differentiation and cell sorting. In addition, we propose a potential alternative mechanism in which cell differentiation, driven by a long-range signal, occurs concurrently with cell sorting. Together, our findings introduce a framework in which fate specification and cell sorting are dynamically coupled during Epi/PrE differentiation.

## Ethics

Not applicable.

## Data accessibility

The code is available on GitHub and will be on Zenodo upon publication.

## Declaration of AI use

Github Copilot was used to increase coding efficiency during development. ChatGPT was used to improve the readability and language in parts of the manuscript.

## Authors’ contributions

S.O.: conceptualisation, implementation, data analysis, figures, writing S.MD.: conceptualisation, writing-review & editing SC.F.: conceptualisation, funding acquisition, project administration, resources, supervision, validation, writing-review & editing

## Acknowledgements

The authors thank India Freund for her drawing of the GRN.

## Appendices

**Figure S1.**
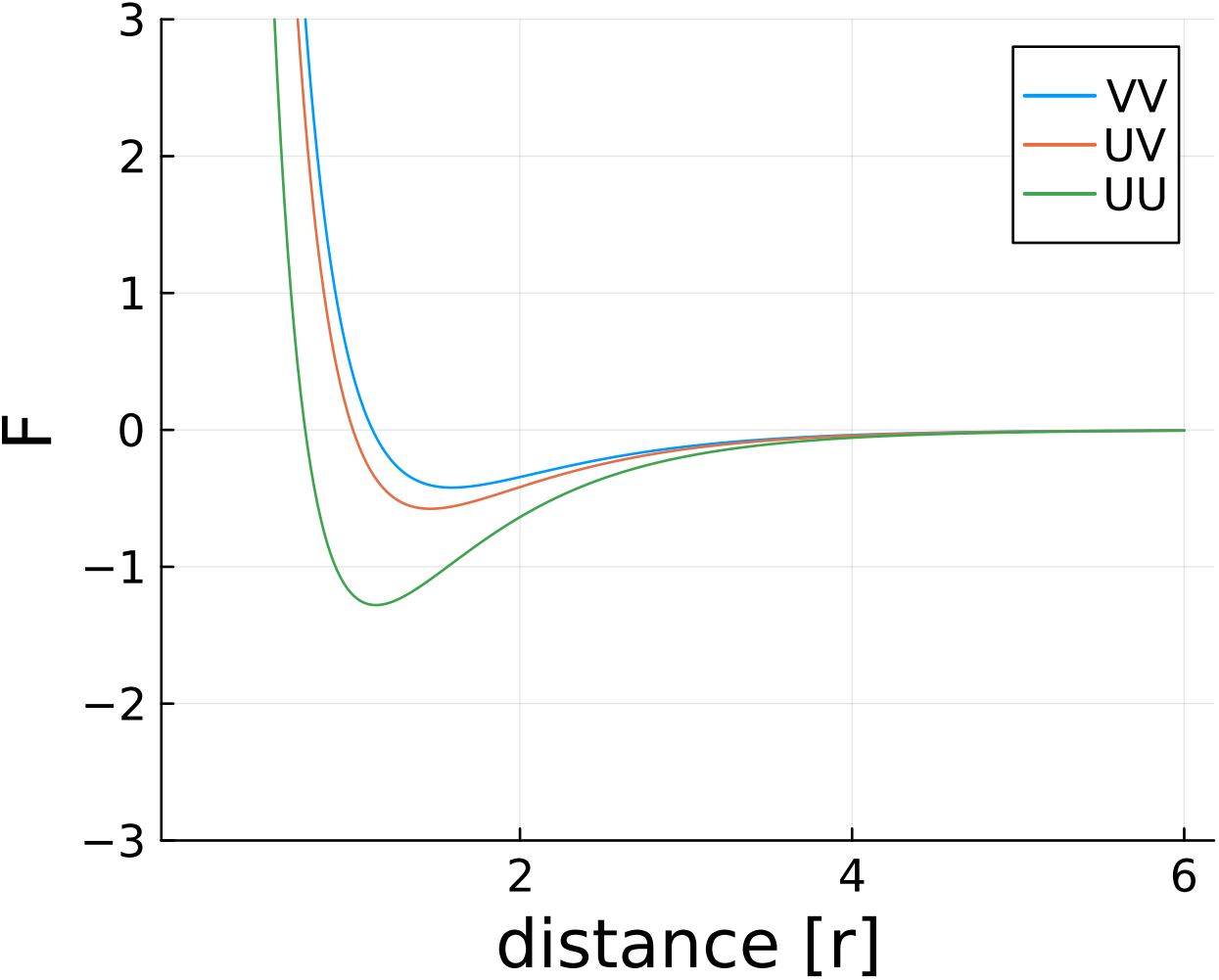
Example forces for the extreme cases of differential adhesion (given in section 2.5) between two cells depending on the distance between them. The distance unit is the radius of the cells.

**Figure S2.**
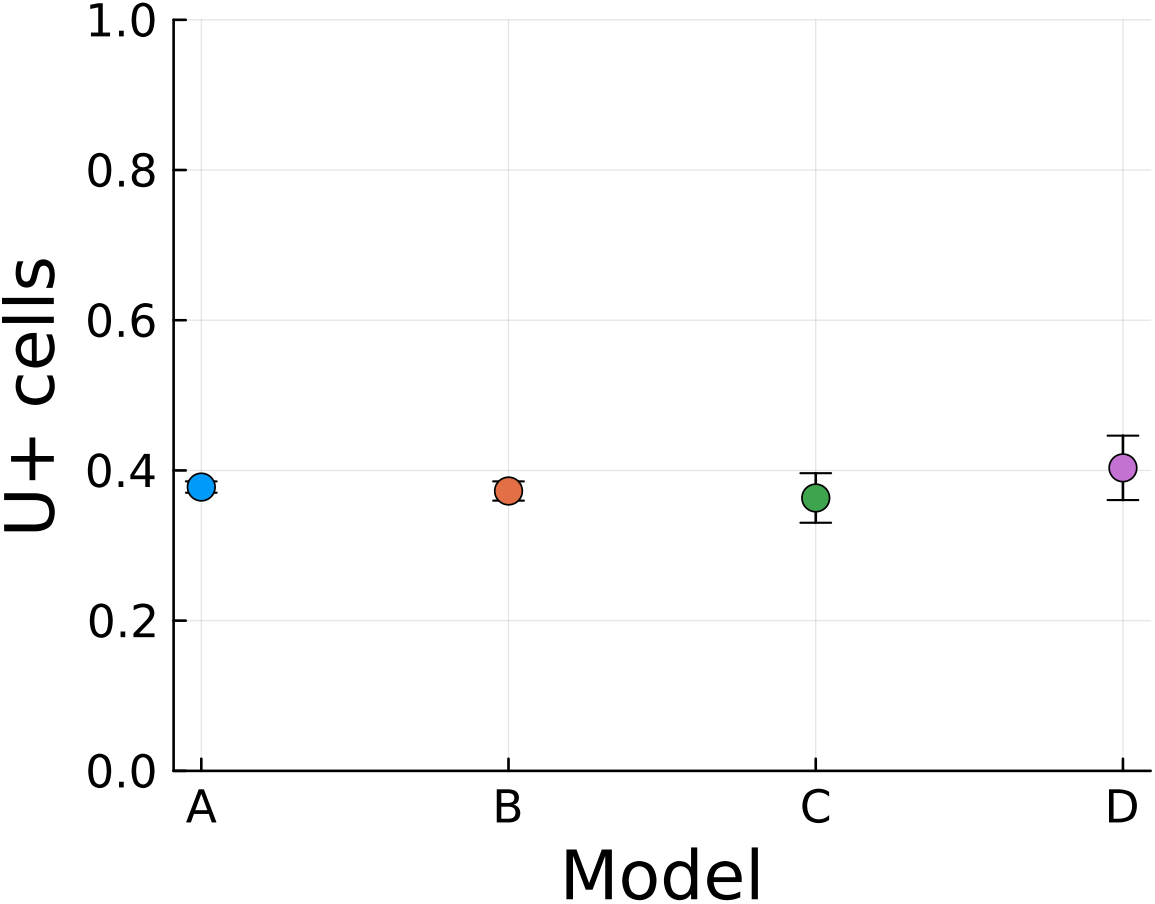
Ratio of U-positive cells at the end of the simulations for all four model variants. Each dot represents the mean for all simulated parameter values of *q* (and *c* for Model D) with standard deviation as the error bars. This leads to a total of 450 simulations for Models A, B and C and 3150 simulations for Model D.

**Figure S3.**
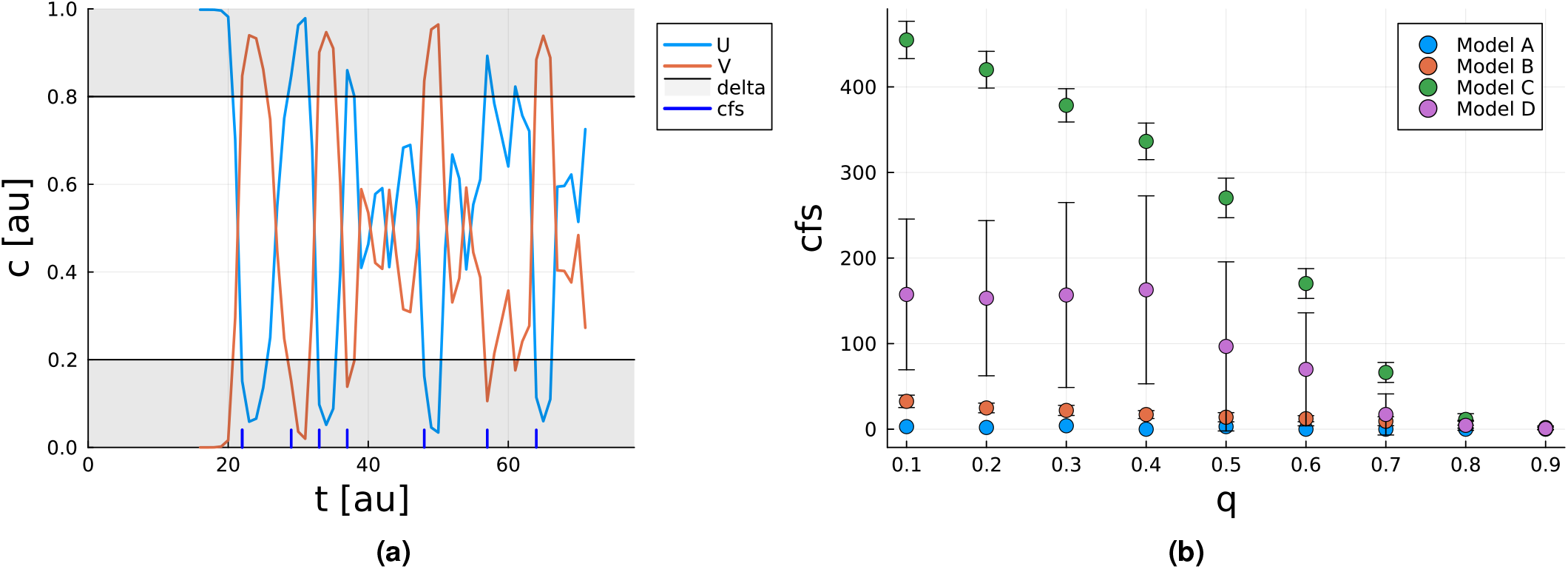
(a) shows the concentration of TFs over time for a single cell after its emergence due to cell division. Blue ticks mark counted cell fate switches. A cell-fate switch is counted when the concentrations of *U* and *V* flip from one grey area 0 + Δ to the other 1 − Δ with Δ = 0.2. **(b)** Mean total cell fate switches (cfs) per simulation for different *q* in all four model versions. Error bars depict standard deviation. For Model D, the same cut-off *c* = 10^−7^ was chosen as for Figure 6 (b). Each dot represents the mean of the total number of cell fate switches across 50 simulations.

**Figure S4.**
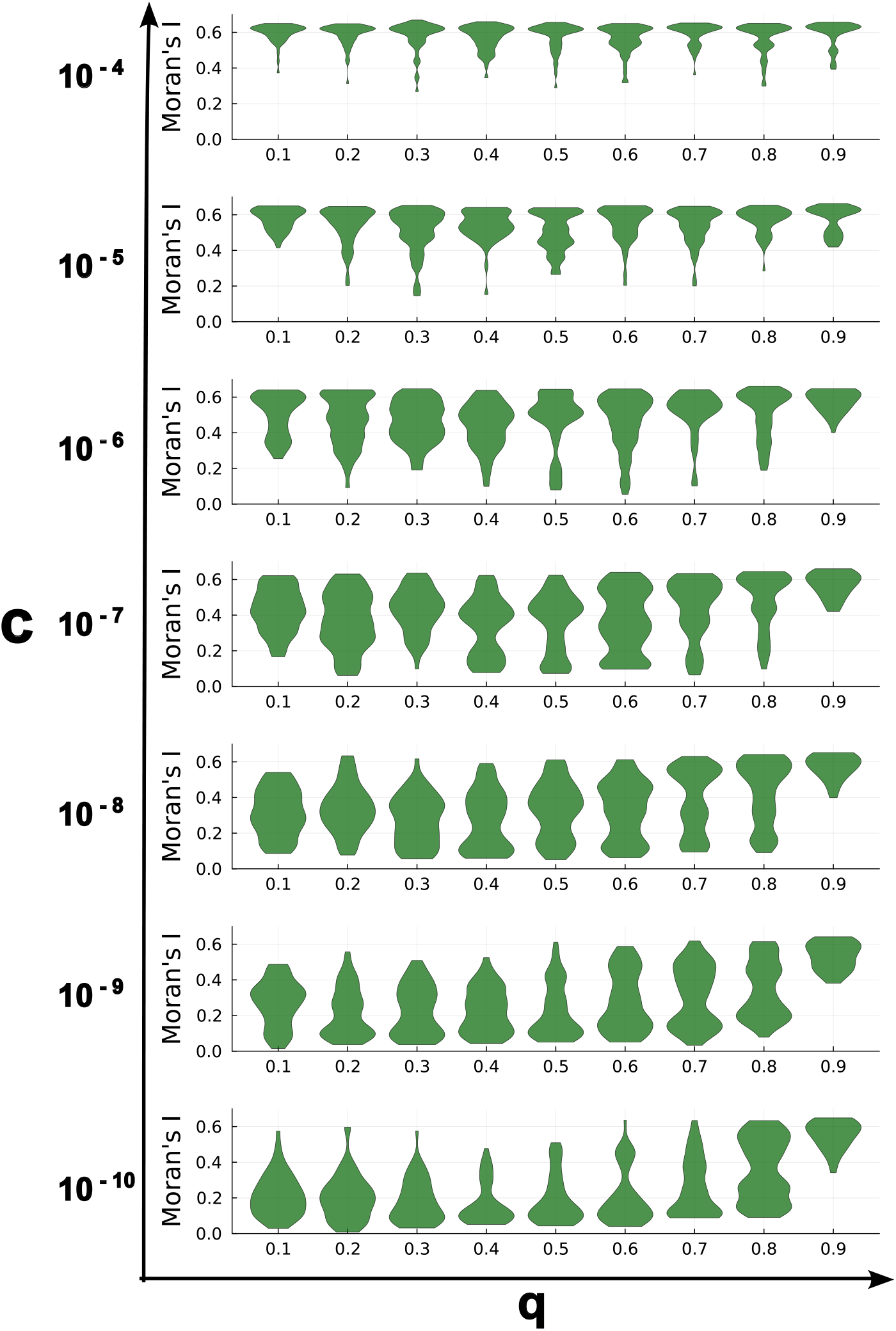
Moran’s I for all simulations used to create Figure 6 (a). Each violin consists of 50 simulations with the shown parameter combination of *c* and *q* for Model D, while other parameter values are given in Table 2. In each run, organoids were simulated until they reached 300 cells.

